# HerbComb: an integrated database for the discovery of novel combinational therapies from herbal medicines

**DOI:** 10.1101/2025.08.06.668811

**Authors:** Yinyin Wang, Rui Liu, Xiang Luo, Jiaqi Yao, Hao Liu, Chengyuan Yue, Wencheng Xu, Xiaochuang Xu, Shixing Lai, Fangcheng Yu, Yinnan Zhang, Lin Cao, Ninghua Tan, Yun Tang, Tao Yang, Jing Tang

## Abstract

Drug combinations have gained increasing interest due to their potential for optimal efficacy and the ability to overcome drug resistance. Herbal medicine is a complex system with multiple ingredients and has illustrated apparent therapeutic effects through thousands of years of clinical experience, offering a rich resource for combinational therapies. However, the underlying mechanism of their synergistic therapeutic effects for disease treatment remains obscure for most herbal medicines. To fill this gap, we developed an integrated database named HerbComb (https://herbcomb.com/) to discover combinatorial therapies in herbal medicine, featuring three unique functions: drug prepositioning, customized combinatorial analysis, and biological and pharmacological features. Additionally, transcriptional gene signatures and ADMET properties were also interrogated as interactive functionalities to facilitate combinational analysis. Using Tongxinluo for Stroke as a case study, we identified and validated Oleanolic acid and Ferulic acid as synergistic ingredients. In summary, HerbComb is a versatile data exploration platform designed to characterize synergistic interactions among herbal medicines, enhancing the understanding of synergistic mechanisms and facilitating the discovery of effective drug combinations.

## 1. Introduction

In recent years, clinicians have increasingly adopted combination therapies to achieve optimal therapeutic effects^1^. Traditional Chinese Medicine (TCM) has demonstrated a unique paradigm of combination therapies, as many herbal formulae employ a multi-component, multi-target, and multi-pathway strategy to yield the intended therapeutic efficacy^2,3^. Although herbal formulae have shown clinical efficacy, the underlying synergistic interactions among their components remain to be fully understood, hindering their broader approval and applications. Although herbal formulae have shown clinical efficacy, the underlying synergistic interactions among their components remain to be fully understood^4-6^, hindering their broader approval and applications.

To harness the therapeutic potential of herbal medicines, advanced computational methods are developed to elucidate their synergistic interactions at the molecular level^7^. In particular, network-based methods have gained increasing popularity for revealing deeper intrinsic connections within these complex relationships^8,9^. Many databases have been developed to support multifaceted network modeling, elucidating the system-level associations between formulas, herbs, ingredients, targets, and diseases. Earlier generations of TCM databases, such as TCM-ID^10^, TCM Database@Taiwan^11^, TCMSP^10^, and TCMID^12^, primarily offered herb-ingredient-target-disease associations with limited modelling capacities. Subsequent advancements led to the construction of more comprehensive network models for TCM, exemplified by TCM-Mesh and an update of TCMID. More recently, databases such as BATMAN-TCM^13^, ETCM^14^, YaTCM^15^, and TCMAnalyzer^16^ have enhanced mechanisms for action analysis. Furthermore, the SymMap database^17^ connected TCM symptoms to modern medical symptoms and diseases. However, most existing databases have focused on characterizing single herbs while ignoring the interactions among multiple herbal components within the same formula^18^. Despite the increasing volume of TCM data, there is currently a lack of integrative tools to explore how these interactions contribute to therapeutic efficacy.

To address this gap, we have developed HerbComb, a comprehensive data integration platform designed to construct an atlas of herbal combinations. HerbComb enables the exploration of herbal combinations at the systems medicine level, which holds significant potential to advance the discovery of novel combination therapies derived from herbal medicine.

## 2. Data collection and modelling

### 2.1 Data extraction and standardization

We extracted all available data from 15 TCM databases. Herb names were standardized into their Pinyin names, and ingredients from different databases were standardized using their InChiKey identifiers. Herbal formulae were collected from six databases, including TCM-ID^13^, TCMID^19^, YaTCM^15^, ETCM^20^, TCMIO^21^, and TCMAnalyzer^16^. Compound-target profiles were extracted from public databases, including ChEMBL^22^, BindingDB^23^, GtopDB^24^, DrugBank^25^, DGiDB^26^, and STITCH^27^. Additionally, we applied a wSDTNBI methodology to predict the targets of ingredients by the structural similarity between herbal ingredients and drugs^28^. Disease-target associations were extracted from a Multi-Scale Interactome (MSI) model^29,30^. Diseases were standardized using the UMLS database^31^, while all the targets were standardized via the UniProt API^32^. 119 ADMET properties were predicted by admetSAR 3.0^33^ for 35,385 herbal ingredients.

We retrieved drug combinational records about herbal medicines from ∼4,000 pieces of literature published between 2000 and 2023, containing the keywords “TCM,” “herbal medicine,” and “combination,” or “synergy.” Herbal perturbation data were retrieved from various resources, including the HERB^34^ and ITCM^35^ databases.

### 2.2 Network-based inference of synergistic ingredients and herbs

In our previous study, we developed a network-based inference of synergistic ingredients and herbs, with rigorous statistical validation, including permutation tests and cross-disease benchmarking^36^, demonstrating that our model can discriminate between herb pairs with high accuracy. Recently, we also applied these methods to understanding the molecular basis of herbal medicines for cough and asthma^37^. Additionally, recent studies have validated the potential of network proximity-based synergy prediction on various diseases^38-40^. The growing consensus supports the validity of our strategy. This motivates our aim to create a user-friendly database with network modelling tools that benefit a broader range of herbal medicine researchers, including those without programming expertise.

Network proximity methods were employed to explore the combinational relationships among prescriptions further, calculating ingredient-ingredient, herb-herb, ingredient-disease, and herb-disease interactions. To explore synergistic ingredients in herbal medicines, the network proximity method^36,41^ as employed to determine an interaction score for an ingredient pair as follows:

#### Ingredients-ingredient synergistic score

For all potential ingredient pairs within a single formula, we applied a closest distance algorithm based on their target network to calculate their combinational distance. The closest distance algorithm is defined as follows:

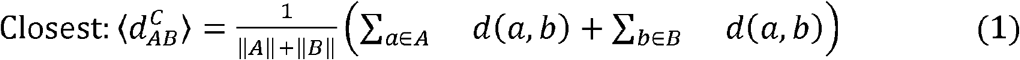

In function 1, *A* and *B* are set of targets separately. *d*_(a,b)_ is the shortest path length between target *a* in network *A* and target *b* in network *B*. ∥ *A* ∥ and ∥ *B* ∥ denote the number of nodes in networks *A* and *B* separately. For each node *a* in network *A*, its shortest path length to each node in herb *B* is calculated and the minimum one will be kept. The exact process is applied to network *B*. Finally, the minimum values of each node in both networks *A* and *B* are summed and averaged, indicating how closely these two networks interact. Her, *A* represents the targets of one ingredient, and *B* represents the targets of another ingredient. Here, smaller *d* (*X, Y*) indicate strong interactions among PPI networks.

#### Herb-herb interaction score

The distance between herb-herb combinations was calculated based on their ingredient-ingredient network, with the shortest distance as the edge weight. The distance between two herbs is the average shortest distance of their center ingredients:

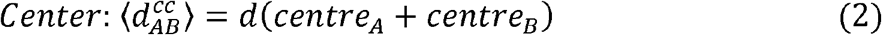

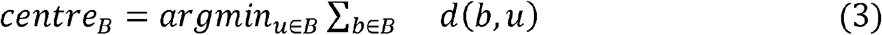

Where *B* is the subnetwork covering all the ingredients in one herb. *d* (*b, u*) representsthe shortest path of each pairwise ingredient within a herb. The central ingredient is the one with the minimum sum distance to other ingredients. The shortest algorithm for calculating the distance between two ingredients is defined as follows:

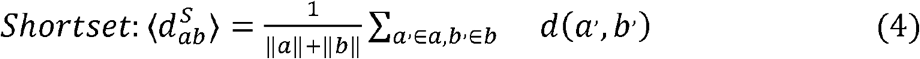

Where *a*, and *b*, are the targets from ingredient *a* and *b* separately. The product of ∥ *a* ∥ and ∥ *b* ∥ represents the number of targets for ingredients *a* and *b*. Finally, the shortest ⟨ *a, b* ⟩ is the sum of all shortest path lengths between the two target sets, averaged to provide a measure of network proximity.

To deteramine the statistical significance, the null distribution was calculated using 1000 random ingredient pairs. Here, we performed a one-tailed Fisher’s Z-test for each pair using the formula: *Z*= (*x* − *µ*_*null*_)/*σ* _*null*_, where *x* is the observed proximity score, *µ*_*null*_ is the mean of the null distribution, and σ _null_ is the standard deviation of the null distribution. Pairs with P < 0.05 were considered statistically significant. Therefore, in this study, we used all co-occurring pairs in formulae as the null distribution, comprising 61,757 herb-herb and 304,992 ingredient-ingredient pairs, due to computational constraints (exhaustive testing of 2.5 billion pairs being infeasible). For disease associations, we generated null distributions from 1,000 randomly sampled pairs per test.

Similarly, the interaction scores between an herb-herb pair are determined based on their ingredient-ingredient network. Specifically, the interaction score is the average shortest distance between union targets of all their ingredients. Herb pairs with the lowest 5% interaction scores are classified as synergistic herb-herb combinations.

### 2.3 Network-based inference of herb disease indications

To further predict the disease indication of herbs, the network proximity method was employed to determine the association scores.

#### Ingredient-disease association score

To measure the association between ingredients and disease, we applied the z-score method. This network module method evaluates the relatedness between one drug and one disease. The shortest path length between drug X and disease Y is defined as:

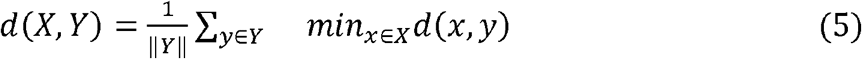

*d*(*x, y*) is the shortest path length between target node *x* in drug *X* and target node *y* in disease *Y*. The minimum values across all nodes in disease *Y* are summed and averaged, similar to the closest distance algorithm but in one direction from *Y* to *X*.

#### Herb-disease association score

Similar to ingredient-disease associations, we combined all the targets of ingredients in one herb as the target set for one herb. Denoting that *c*_*hA*_ = (*c*_*1*_, *c*_*2*_,…, *ci*) is a set ingredient in herb *A. c*_*hB*_ = (*c*_*1*_, *c*_*2*_,…, *ci*) is a set ingredient in an herb *B*.*T*_*C*_ = (*T*_1_, *T*_2_,…,) is a set of targets in one compound. Then, the target for one herb can be denoted as:

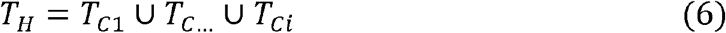

*T*_*H*_ is the union of targets of ingredients in this herb *X*. We extracted disease-related genes *Y*. The herb-disease proximity was also calculated.

For ingredient-disease associations, the shortest path length *d* (*X, Y*) between drug *X* and disease *Y* is defined:

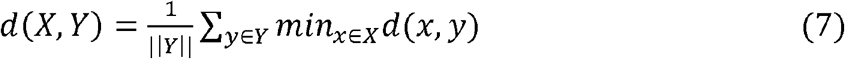

Here, *d*_(*x,y*)_ represents the shortest path length between a target node x in drug X and target node *y* in disease *Y*. The null distribution of *d*(*X, Y*) was determined by 1,000 random drug-disease pairs, based on which a *P* < 0.05 was considered significant. Similarly, for herb-disease associations, we combined all the targets of the ingredients within herbs to form a target set for that herb.

### 2.4 Technical Implementation

HerbComb is implemented as a web-based platform with a client-server architecture. The frontend is built with Vue.js and Element UI, supporting both English and Chinese through Vue I18n. The backend utilizes Java and Spring Boot frameworks to deliver RESTful APIs and real-time services. Data is stored in MySQL (version 8.0) and cached in Redis (version 6.0). The web server runs on nginx (version 1.18) on TencentOS 5.4. Data processing pipelines were developed in Python (using Pandas and NetworkX) to integrate and clean data from multiple sources. Core entity data (diseases/conditions, herbs, ingredients, and targets) is available as downloadable CSV and JSON files from the website. In contrast, association data (herb-target, ingredient-target, herb-herb) can be exported per query through the interface. All data is released under a CC BY 4.0 license, permitting free download, reuse, modification, and redistribution, as well as commercial use.

## 3. Results

### 3.1 Overview of HerbComb database

HerbComb was designed to facilitate the identification of synergistic interactions among herbal components. To achieve this goal, various available information on TCM was integrated into the **HerbComb** database, encompassing 46,929 formulas, 7,516 herbs, and 35,385 ingredients. Additionally, to provide a comprehensive understanding of the mechanisms of action and disease indications, the database includes 25,212 disease-associated genes, 162,299 herb-ingredient pairs, 348,543 ingredient-target pairs, and 325,212 disease-target pairs, all organized into a multi-level heterogeneous network (**Figure 1** and **Figure 2B-D**). Based on these prior acknowledgments, we employed network-based methodologies to systematically characterize interactions among ingredients, thereby constructing a comprehensive combinatorial landscape for herbal medicine (**Figure 1** and **Figure 2A**).

**Figure 1.**
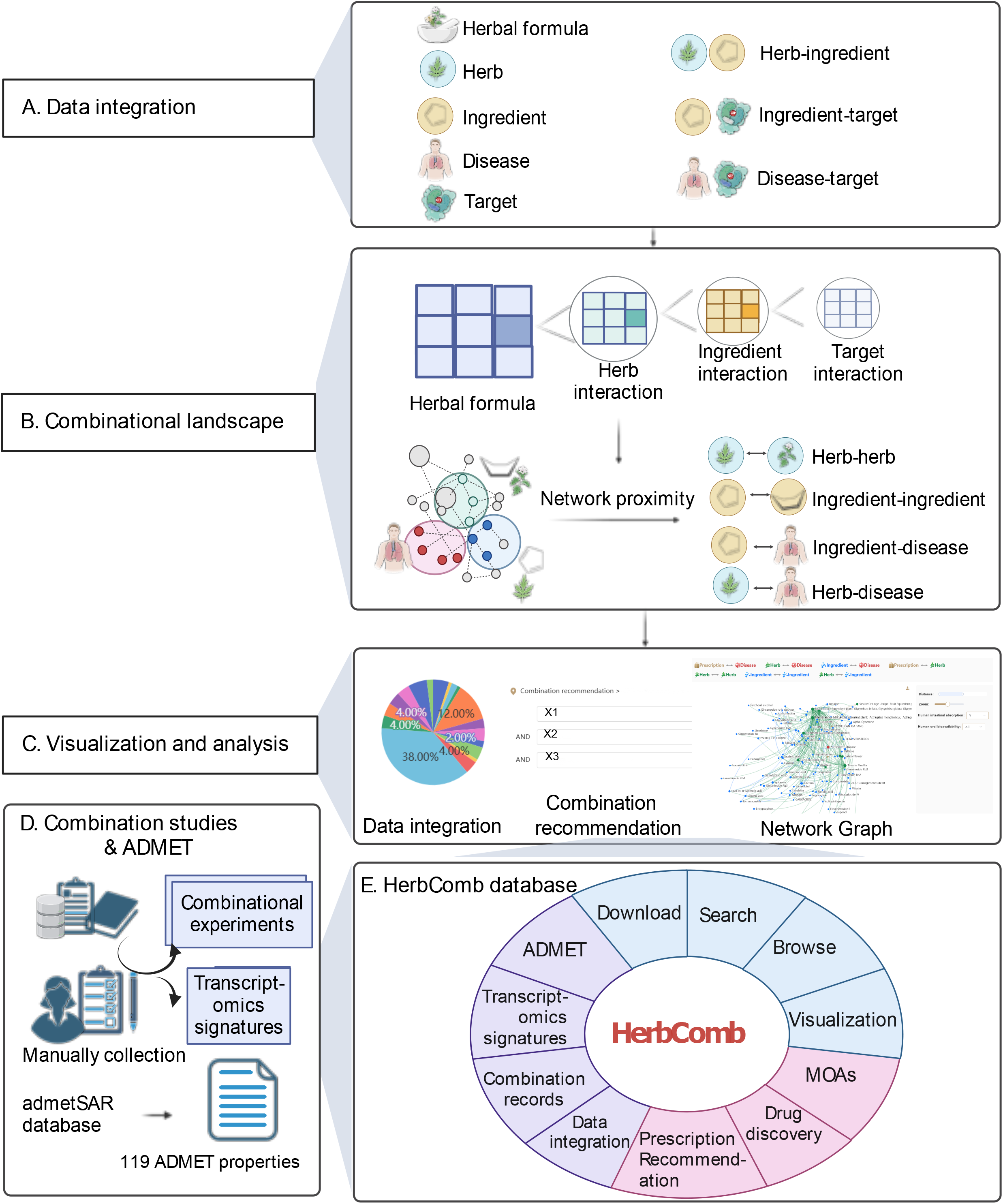
The overview of the HerbComb database. **(A)** HerbComb integrates formula-herb-ingredient-target associations, protein-protein interactions, and disease-gene associations, covering 851,409 ingredient-disease pairs (839 diseases and 1,016 ingredients), 3,053,258 herb-disease pairs (838 diseases and 3,644 herbs), 304,992 ingredient-ingredient pairs and 61,757 herb-herb pairs that co-occur in 46,929 TCM formulae. **(B)** A network proximity model was constructed to characterize the interactions between herbs, diseases, ingredients, and their combinations. Downstream analysis of the synergistic interactions was provided, effectively capturing the unique combinatorial characteristics of herbal medicines. **(C)** Customized combinational analysis: HerbComb supports tailored analyses between ingredients, herbs, TCM prescriptions, and diseases through the “Combination Recommendation” button. This function mimics the prescription process in TCM clinical practice. **(D)** Additionally, experimental data about herbal combinations and transcriptomics signatures were manually curated from the literature, and ADMET properties were systematically calculated. **(E)** HerbComb is a versatile data exploration platform designed to characterize synergistic interactions among herbal medicines, enhancing the understanding of synergistic mechanisms and facilitating the discovery of effective drug combinations for disease treatment.

**Figure 2.**
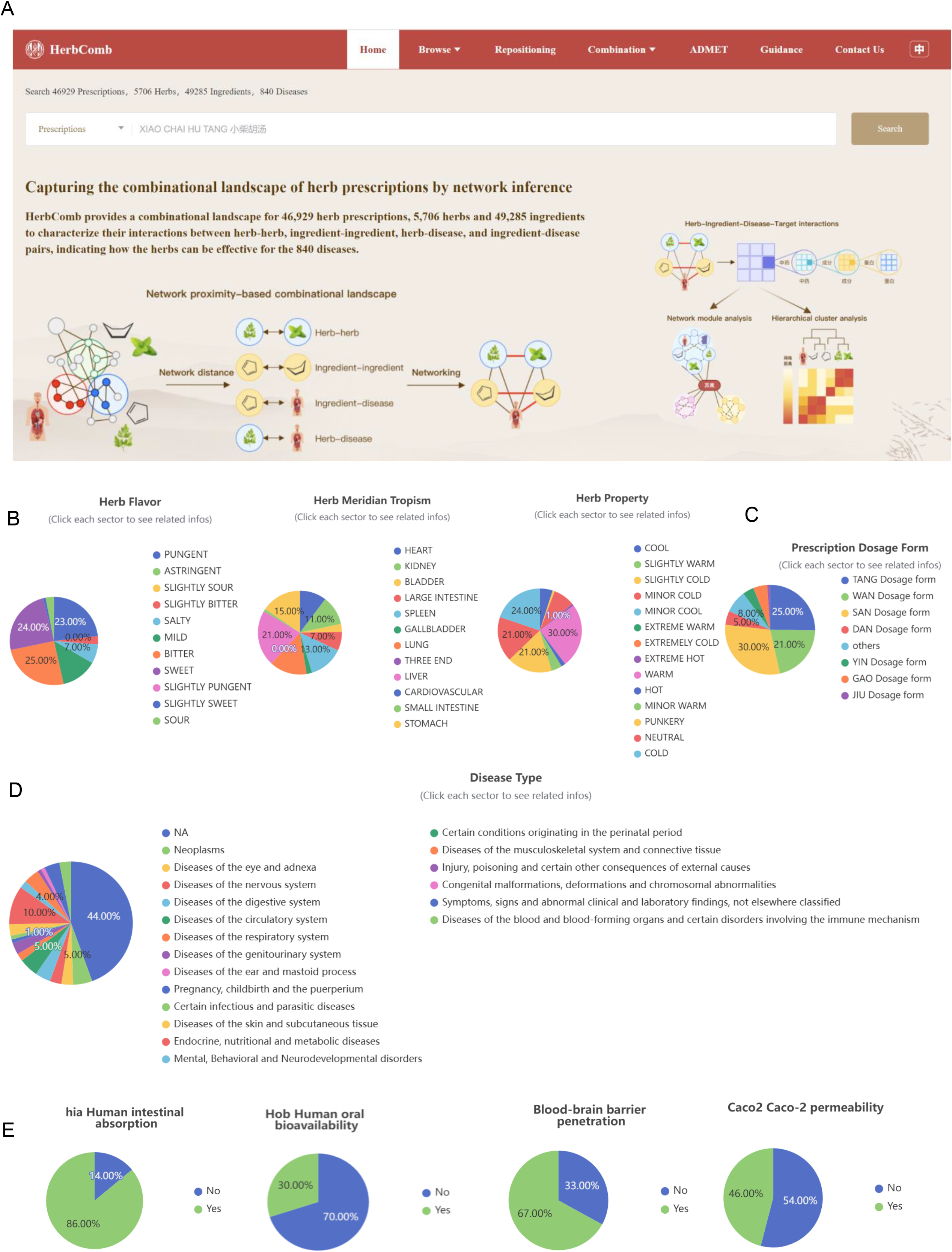
Data statistics of HerbComb. **(A)** Home page of HerbComb. **(B)** Distribution of flavors, meridians, and properties for herbs in HerbComb. **(C)** Distribution of dosage forms of prescriptions in HerbComb. **(D)** Distribution of disease types contained in HerbComb. **(E)** Distribution of ADMET properties for ingredients in HerbComb.

HerbComb offers the following four innovative functionalities (**Figure 1**), including: (i) the identification of potential therapeutic and synergistic herbal components for a wide range of diseases; (ii) interactive analysis of the underlying mechanisms of action for specific herbal formulations; (iii) a decision-support tool to predict the therapeutic effects of a given herbal formula, emulating the prescription process of TCM; and (iv) the integration of pharmacotranscriptomics, pharmacokinetics, and toxicity profiles, facilitating a systems-level characterization of herbal medicine.

### 3.2 Screen therapeutic herbs and ingredients with synergistic effects

HerbComb employs systematic modeling to identify potential therapeutic and synergistic ingredients or herbs for specific diseases^42^. By clicking the “Repositioning” button, users can browse all potential therapeutic herbs and ingredients for 840 diseases (**Figure S1**).

Specifically, we constructed a substantial combinational atlas, covering 851,409 ingredient-disease pairs (839 diseases and 1,016 ingredients), 3,053,258 herb-disease pairs (838 diseases and 3,644 herbs), 304,992 ingredient-ingredient pairs, and 61,757 herb-herb pairs that co-occur in 46,929 TCM formulae. Consequently, HerbComb identified 2,999 high-confidence synergistic herb-herb pairs, 7,748 ingredient-ingredient interactions, 179,461 therapeutic disease-herb associations, and 27,673 disease-ingredient associations across 840 diseases. This extensive resource significantly complements existing databases (**Figure 3A)**.

**Figure 3.**
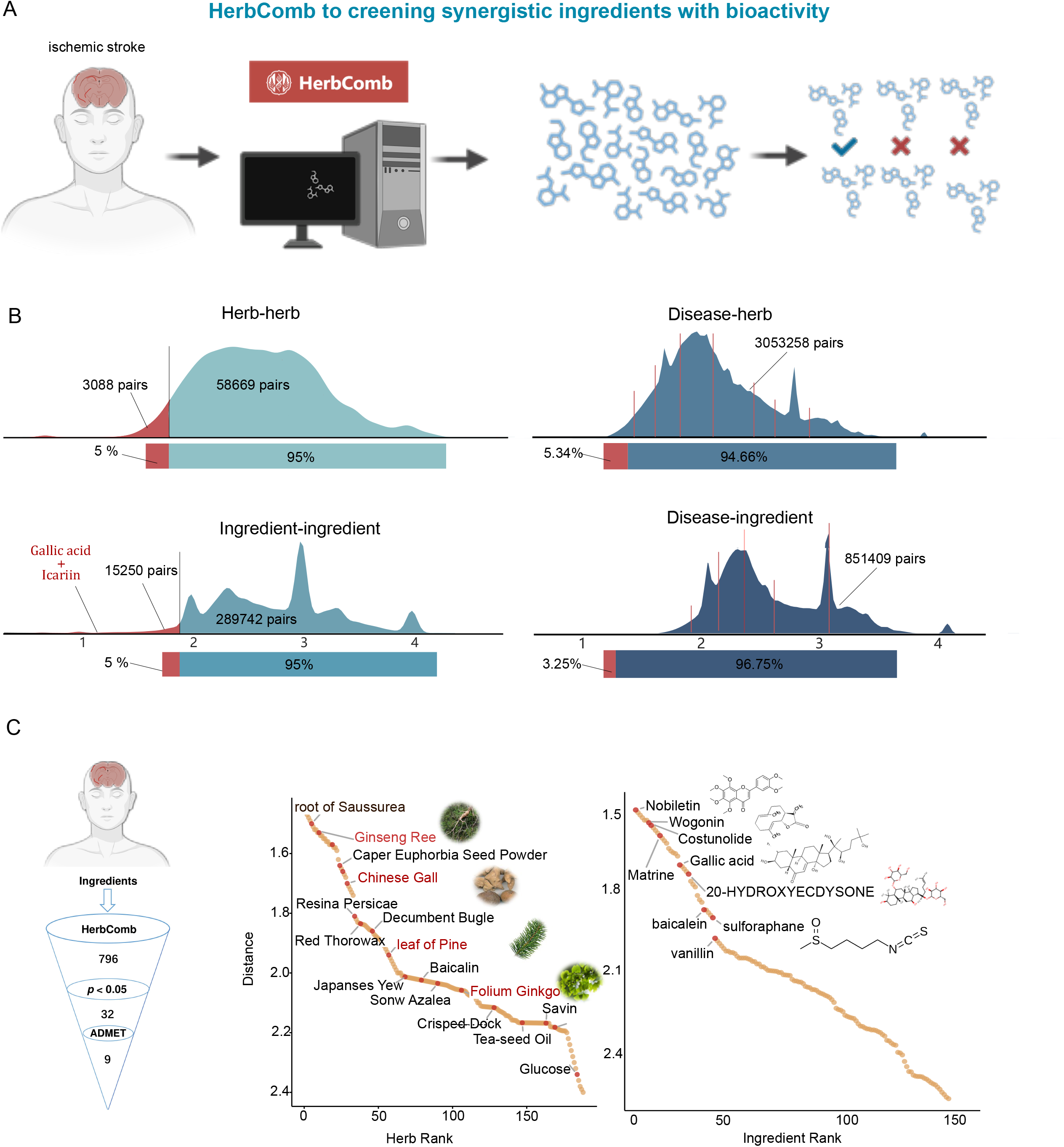
Large-Scale screening results of therapeutic herbs or ingredients for 840 Diseases. **(A)** Concepts of drug repurposing function: HerbComb introduces a drug repurposing feature to identify potential therapeutic applications of herbs and ingredients across a wide range of diseases. **(B)** Statistical analysis of significant associations: The HerbComb platform offers statistical insights into the significant relationships between herb-herb, herb-disease, ingredient-ingredient, and ingredient-disease pairs, facilitating a deeper understanding of their interactions. **(C)** Drug screening for ischemic stroke: Using ischemic stroke as an example, HerbComb screens and prioritizes top therapeutic herbs and ingredients, which are then visualized for further analysis.

On average, four herbs and one ingredient can be associated with each disease (**Figure 3B**), highlighting the potential of herbal medicines as a valuable resource for drug discovery. Additionally, HerbComb serves as a comprehensive data portal for uncovering therapeutic and synergistic ingredients within TCM formulas. On average, six herbs and 199 ingredients in each herbal formula are significantly associated with specific diseases.

Neurological disorder Stroke is an area where herbal medicines have demonstrated promise^43,44^. Using the “Repositioning” function in **HerbComb**, we screened potential herbal medicines and ingredients for stroke as an example. Consequently, 32 ingredients were prioritized with statistical significance (P < 0.05) with ischemic stroke (**Figure 3C**). Furthermore, when searching for a particular herb or ingredient, the top-scoring diseases will be provided as potential novel indications on which that herb/ingredient may have therapeutic efficiency. Among these repurposed herbs, *Ginseng Reed, Chinese Gall, leaf of Pine*, and *Folium Ginkgo* have been reported to be associated with ischemic stroke, underscoring the potential of HerbComb for drug discovery. Similarly, prioritized ingredients, such as Ginsenoside RG1^45^, Ginsenoside Rg3, ferulic acid, and Cryptotansl, have also been reported for ischemic stroke treatment.

### 3.3 HerbComb offers a comprehensive platform for combinatorial analysis

1. HerbComb offers a comprehensive platform for combinatorial analysis, providing interaction scores for millions of herb and ingredient pairings (**Figure 1B**). The platform features six main functions designed to facilitate in-depth exploration and analysis:
2. Herb-specific synergistic interactions: When users search for a specific herb for or prescription, HerbComb returns significant synergistic ingredient-ingredient interactions within that herb or prescription.
3. Herb-Herb combinations and disease associations: The HerbComb platform provides all potential herb-herb combinations and diseases associated with the searched herb.
4. Ingredient-Ingredient combinations and disease associations: HerbComb provides all potential ingredient-ingredient combinations and diseases associated with the searched ingredient.
5. Visualization of synergistic pairs: These significantly associated pairs can be visualized in a comprehensive combination network, enabling downstream analyses such as modularity and ADMET filtering.
6. Customized combinational analysis: HerbComb supports tailored analyses between ingredients, herbs, TCM prescriptions, and diseases through the “Combinational Recommendation” button. This function mimics the prescription process in TCM clinical practice. In total, five combination patterns are supported in the “Combinational Recommendation” function (**Figure 4A**): “herb + herb + disease,” “ingredient + ingredient + disease,” “prescription + prescription + disease,” “herb + herb,” and “ingredient + ingredient.” For example, users can query the therapeutic potential of a specific herb combination for treating a particular disease (i.e., the “herb + herb + disease” query pattern). After inputting the herbs and disease, a combinatorial landscape is visualized, highlighting interactions among herb-herb, ingredient-ingredient, herb-disease, and ingredient-disease pairs (**Figure 1C**).

**Figure 4.**
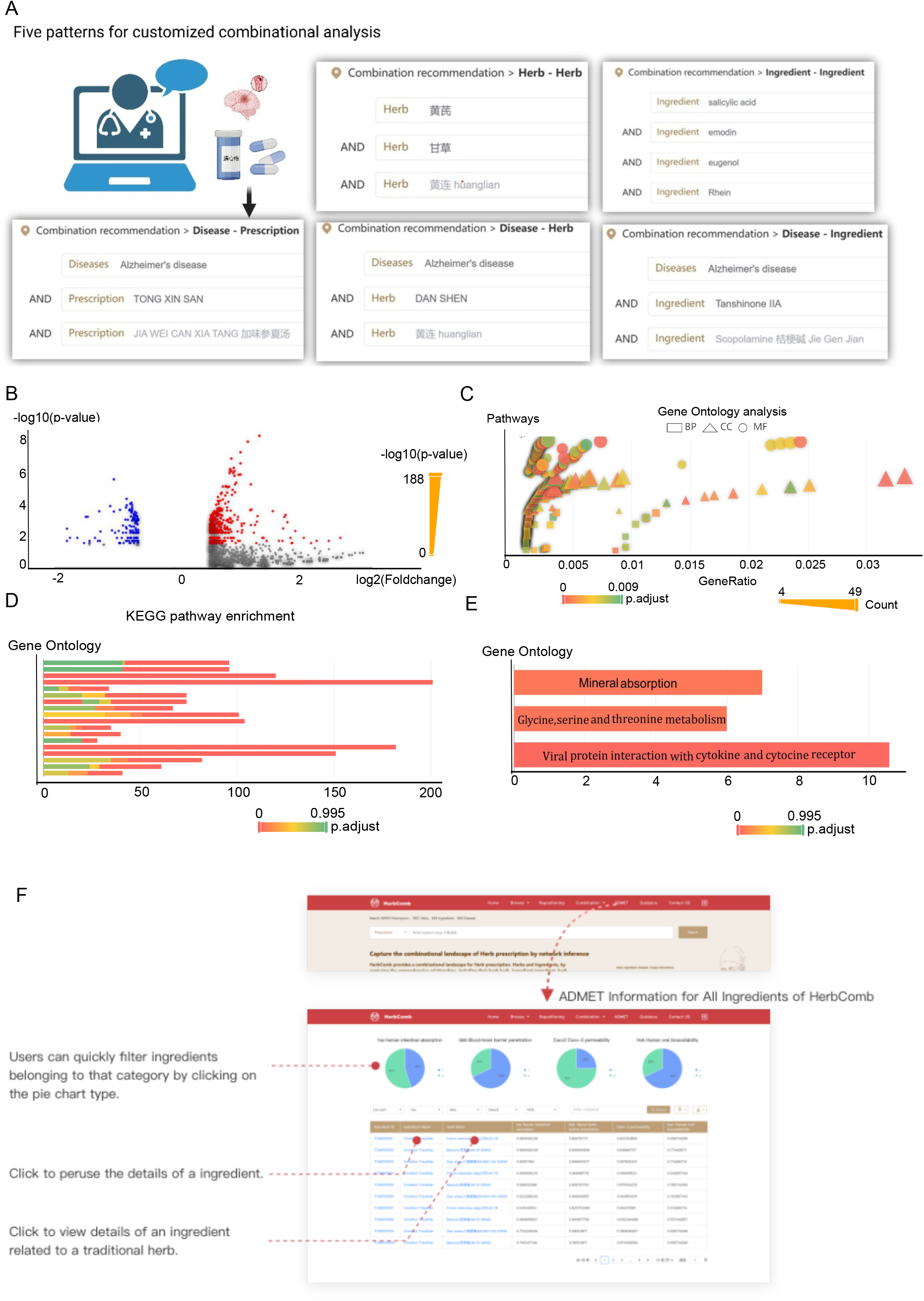
Additional functions of HerbComb to enhance combinational analysis. **(A)** HerbComb enables tailored analyses between ingredients, herbs, prescriptions, and diseases through the “Combination Recommendation” feature. This function supports five combination patterns: **a)** “herb + herb + disease”, **b)** “ingredient + ingredient + disease”, **c)** “prescription + prescription + disease”, **d)** “herb + herb”, and **e)** “ingredient + ingredient”. (B-D) Transcriptomics signature analysis for determining synergistic mechanisms. HerbComb integrates transcriptomics data to uncover the molecular mechanisms underlying synergistic interactions, providing deeper insights into how herb and ingredient combinations exert their effects. **(E)** Enriched pathways of Ginsenoside Rg1 in the treatment of stroke by transcriptomics signature analysis. **(F)** HerbComb enables the systematic evaluation of ADMET (Absorption, Distribution, Metabolism, Excretion, and Toxicity) properties for ingredients in herbal medicines, facilitating more efficient drug screening processes.

In addition to predicted combinations from network modeling, HerbComb provides curated data on experimentally validated herbal or ingredient combinations. Systematically curating around 4,000 publications offers comprehensive information on 79 herb-drug, 98 ingredient-drug, 84 herb-herb, 76 ingredient-ingredient, and 14 ingredient-herb interactions, filling a critical gap and serving as a foundational resource for computational models.

HerbComb is particularly valuable for experimentally validated combinations whose molecular synergistic mechanisms remain unclear as it supports in-depth combinatorial landscape analysis, enhancing our understanding of known synergistic pairs. For example, Icariin and Gallic acid have been reported to exhibit synergistic effects in Alzheimer’s disease^46^. Using the “Combinational Recommendation” function, HerbComb revealed that these two compounds exhibit stronger interactions and are significantly associated with Alzheimer’s disease (**Figure 3B**). As shown in **Figure 3B-C**, the combination of Icariin and Gallic acid ranks 2306 across all pairs. Additionally, the distance between Icariin and AD is 2.03 (*P* = 0.03), while the distance between Gallic acid and AD is 1.97 (*P* = 0.03). This suggests that Icariin and Gallic acid may have synergistic therapeutic effects. While our model identifies statistically significant compound-disease associations through network proximity, these represent potential mechanistic relationships rather than confirmed treatment effects. Clinical validation remains essential, particularly since network proximity may capture both therapeutic and pathophysiological associations. It was reported that *Epimedium* aqueous extract,which contains Icariin as a major active constituent, ameliorates cerebral ischemia/reperfusion injury through inhibiting ROS/NLRP3-mediated pyroptosis^47^. Moreover, Rosemary (*Rosmarinus officinalis* L.), which contains Gallic acid, exhibits neuroprotective effects by enhancing cerebral ischemic tolerance in experimental stroke^48^. Similarly, green tea, a natural product rich in Gallic acid, also shows potential therapeutic effects against ischemic stroke^49^. This finding highlights the potential of HerbComb to uncover and elucidate the mechanisms of action (MOA) underlying experimental combinatorial results.

### 3.4 Transcriptomics signature analysis for synergistic mechanisms determination

Typically, target information for most herbs is predicted using computational tools but often lacks experimental validation, which can introduce noise. Recently, transcriptomic changes induced by herbal medicines, referred to as gene expression signatures, have shown promise in elucidating the holistic effects of specific herbs or ingredients^50,51^. Therefore, the HerbComb database provides gene expression changes induced by herb perturbations before and after treatment, establishing differentially expressed genes (DEGs) as transcriptomic signatures (**Figure 1D**). These signatures enable the identification of molecular pathways influenced by the treatment, facilitating a deeper understanding of the pharmacological mechanisms underlying herbal therapies.

HerbComb has now integrated data from various herbal perturbations to generate gene expression signatures for 693 herbal treatments. It also offers tools to analyze and visualize DEGs, pathway enrichment, and Gene Ontology (GO) terms (**Figure 4B-D**), thereby enhancing the understanding of transcript-level effects of herb or ingredient perturbations. These gene signatures serve as a critical complementary dataset for analyzing herbal combinations, aiding in identifying synergistic mechanisms^52^.

For instance, although Ginsenoside Rg1 has been prioritized for ischemic stroke by the repositioning function of HerbComb (**Figure 3C**), the underlying therapeutic mechanism remains unknown. Therefore, transcriptomics signature analysis was also supported by HerbComb to facilitate further exploration of synergistic mechanisms. In total, 24 signature genes were associated with Ginsenoside Rg1 in HerbComb, which were mainly enriched mineral absorption, glycine, serine, and threonine metabolism, and viral protein interaction with cytokine and cytokine receptors (P < 0.05, **Figure 4E**). Intriguingly, Ginsenoside Rg1 has been reported to regulate cytokines through the viral protein interaction with cytokine and cytokine receptor pathway^53^, thereby underscoring HerbComb’s potential to provide mechanistic insights into herbal treatments.

### 3.5 ADMET properties to facilitate the identification of synergistic ingredient

Studying ADMET properties is crucial for understanding the pharmacokinetics and pharmacodynamics of herbal medicine. While existing databases provide some ADMET properties (e.g., TCMSP, YaTCM, ETCM, and TCMID), they often lack critical information, particularly on toxicity. HerbComb addresses this gap by integrating the most up-to-date set of 119 ADMET properties for 35,385 ingredients, encompassing 18 physicochemical properties, 43 human health toxicity endpoints, 16 environmental risk assessment endpoints, and nine cosmetic risk assessment endpoints (**Figures 1D** and **4F**). This comprehensive ADMET information is invaluable for identifying synergistic ingredients with high drug-likeness and low toxicity.

As shown in **Figure 2E**, approximately 86% of the 9,783 ingredients exhibit optimal human intestinal absorption, while only 30% demonstrate good oral bioavailability. Furthermore, around 67% of the ingredients can pass the blood-brain barrier. HerbComb also provided a list of 18,314 ingredients (36.55%) that may exhibit hepatotoxic effects for an alert.

HerbComb enables users to manually filter ingredients based on ADMET properties when conducting combinational analysis, thereby facilitating the process. For instance, among the 32 ingredients prioritized for ischemic stroke, only nine ingredients are significant for ischemic stroke and have good human intestinal absorption and human oral bioavailability > 30% with permanent BBB (**Figure 3C**).

### 3.6. Sensitivity analysis

To gain a deep understanding of the entire database, we conducted a distance distribution analysis to identify popular herbs and ingredients, as well as diseases that are overrepresented.

We conducted a sensitivity analysis to investigate the distribution of ingredients, diseases, and herbs, identifying hub herbs with a significantly larger number of pairs than others. As shown in Figures 5A-B below, the distribution of herbs, ingredients, diseases, and their associated pairs is normalized. On average, an herbal formula contains six herbs and 199 ingredients that can be significantly associated with certain diseases. These results suggest that herbal medicines may be valuable resources for drug discovery. Notably, we found that the top 10 hub herbs and ingredients are associated with more than ∼300 and ∼200 diseases. Similarly, the top 10 hub diseases are associated with ∼1,000 herbs and ∼200 ingredients separately (**Figure 5A**). On average, each disease is associated with 208 and 32.63 herbs and ingredients, while each herb and each ingredient are associated with 47.30 and 35.39 diseases, respectively (**Figure 5B**). Diseases (n = 840) were classified into 19 therapeutic categories. **Figure 5C-D** shows the distribution of significant herb-disease pairs and significant ingredient-disease pairs across these categories. All pairs of disease-herbs and disease-ingredients are unique, with deduplicated pairs removed. The number of herb-disease pairs per category ranges from 197 to 26,948, with diseases in the Endocrine, Nutritional, and Metabolic classification having the highest representation, at 26,948 related disease-herb pairs. These findings suggest that HerbComb is a valuable data portal for revealing the therapeutic and synergistic ingredients in the TCM formula.

**Figure 5.**
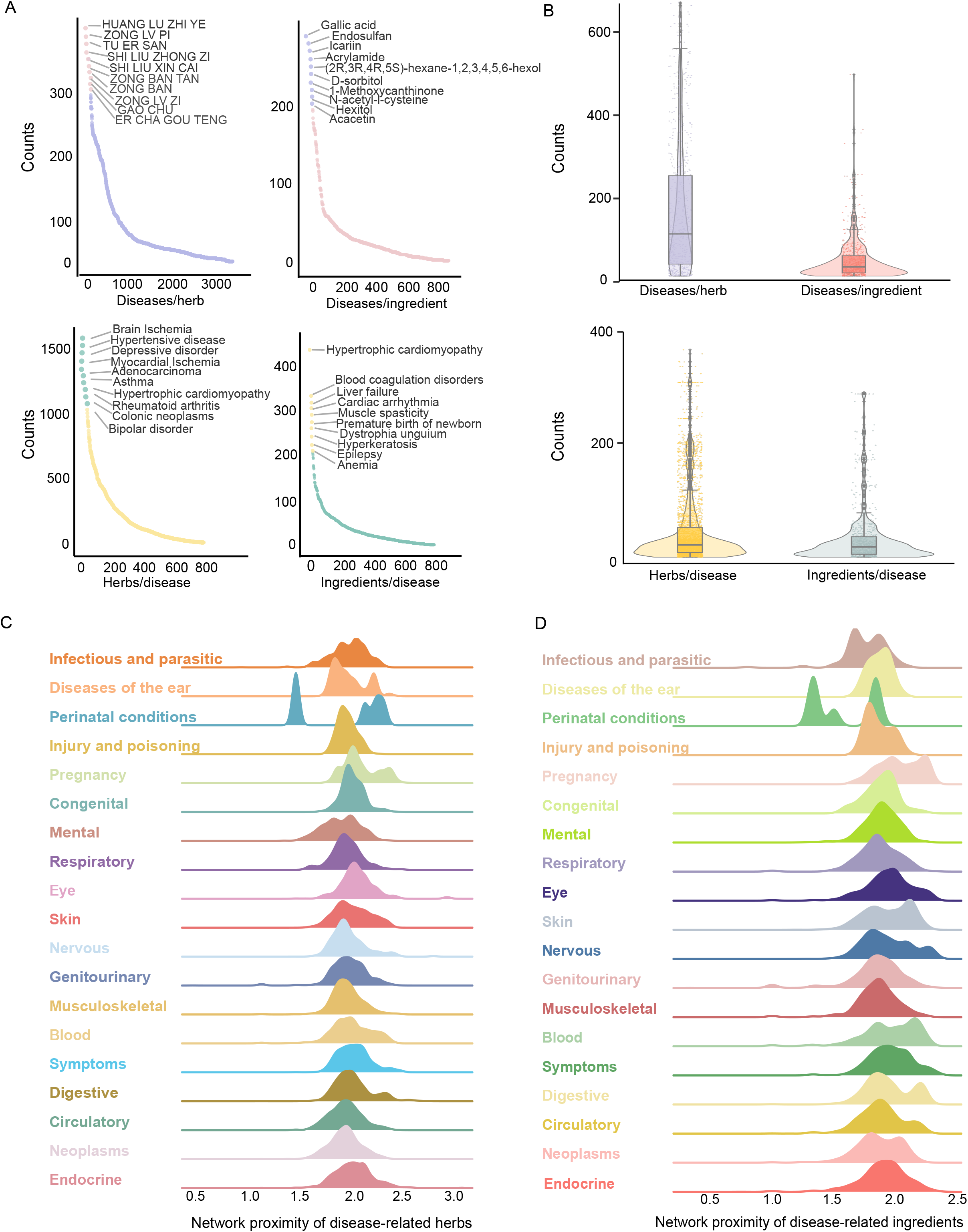
Distance distribution analysis. **(A-B)** Distribution of the number of associated diseases per herb, diseases per ingredient, herbs per disease, and ingredients per disease. **(C-D)** The density of network proximity values for significant herb-disease associations **(C)** and ingredient-disease associations **(D)** across 19 therapeutic classes of diseases.

### 3.7. Stratified analysis

Additionally, we performed stratified analysis by dividing the dataset into meaningful subgroups (called strata), such as herb popularity, disease category, and formula complexity. Stratified analysis helps detect potential differences or biases that might have been masked in the aggregated data.

Firstly, we classified herbs into high- and low-frequency herbs based on their median value in the formulae. We found that higher-frequency herbs show a similar distance to those of lower-frequency herbs (**Figure 6A**). Then, we divided the formula into two groups based on the herbs that comprise it: simple prescriptions with N(herbs) < 6 and complex prescriptions with N(herbs) ≥ 6. Notably, complex prescriptions show significantly closer distances than those simple ones (**Figure 6B**). To characterize the relationship between herb/ingredient and different disease types, we analyzed the distribution of network distances across 19 disease classifications. The bar plot alongside illustrates the number of diseases contained within each category. We observed a similar distribution of distance, regardless of the number of diseases in the classification (**Figure 6C-D**).

**Figure 6.**
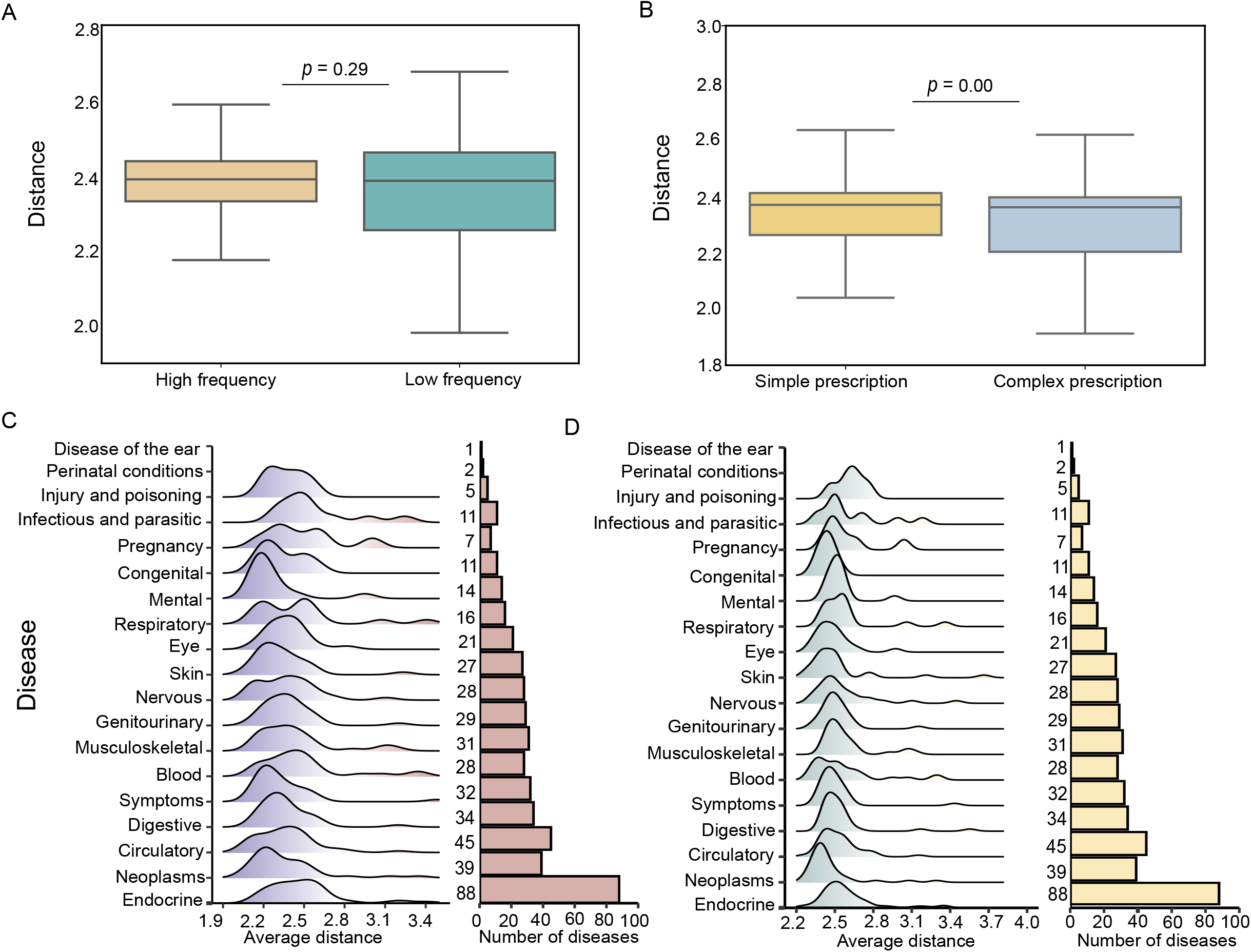
Stratified analysis of distance. **(A)** Distance comparison by classifying herbs into high and low frequency herbs by their median value in formulae. **(B)** Distance comparison by cutting the formula into two groups according to the herbs that comprise that formula: Simple prescription with N(herbs) <6 and complex prescription with N(herbs) ≥ 6. **(C-D)** Network distance distributions between herbs/ingredients and diseases across 19 disease classifications. Ridge plots display distance distributions, while bar plots indicate the number of diseases in each category.

### 3.8. Case study

Using Stroke as a case study, we have identified a famous herbal formula, Tongxinluo, in which we have identified Oleanolic acid and Ferulic acid as synergistic ingredients that have not been previously reported.

The TCM formula Tongxinluo Capsule (TXL) has been used clinically for the prevention and treatment of cardiovascular diseases, particularly ischemic stroke. However, existing research primarily focuses on its overall efficacy, with limited knowledge of the interactions between its ingredients. Using HerbComb, we determined the interaction distances between herbs and ingredients associated with ischemic stroke.

The combinational effects of TXL were further investigated by constructing a network to represent the interactions between ingredients and diseases (**Figure 7A**). To identify key active components and potential synergistic ingredients, the Louvain algorithm was employed for community detection, based on the assumption that ingredients clustered with the ischemic stroke node are more likely to show synergistic effects. According to ingredient-disease and ingredient-ingredient distance analysis, Oleanolic acid and Ferulic acid were ultimately selected (**Figure 7B**).

**Figure 7.**
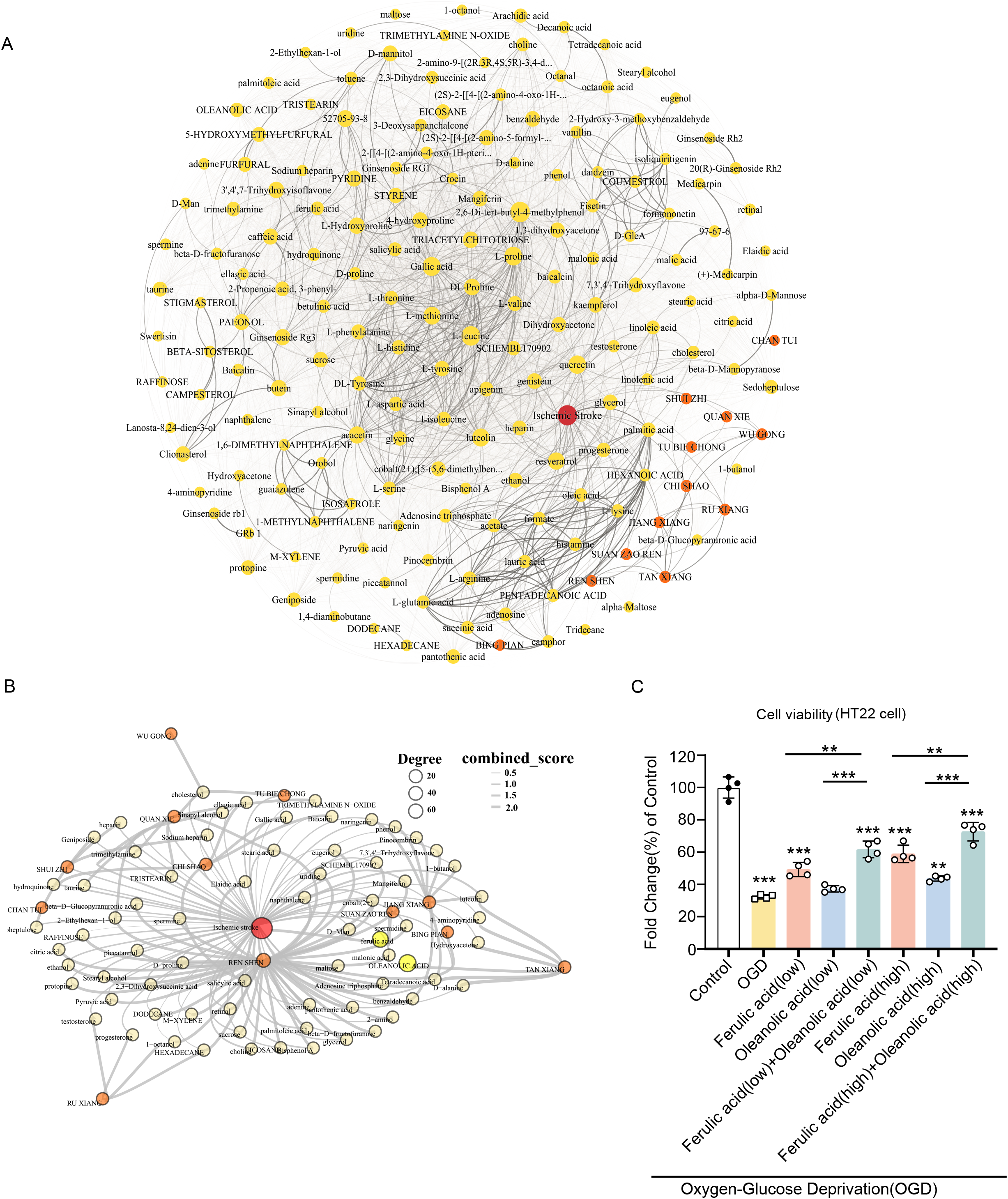
The combinatorial landscape of Tongxinluo Capsule. **(A)** Herb-Component-Disease Network. **(B)** Key disease-associated module via network community analysis. **(C)** The viability of HT22 cells in Oxygen and Glucose Deprivation (OGD) conditions of different concentrations of Oleanolic acid and Ferulic acid (T-test, ^*^ P < 0.05, ^***^ P < 0.01, ^***^ P < 0.001).

To validate the synergistic effects of Oleanolic acid and Ferulic acid, we conducted a cell viability assay on neural protection using the HT22 cell line (immortalized mouse hippocampal neuronal cells). We found that the combination of Oleanolic acid and Ferulic acid at both low and high concentrations shows significantly increased cell viability in the HT22 cell line (T-test, *P* < 0.01, **Figure 7C**). These results suggest that the HerbComb database can be an effective platform for discovering synergistic ingredients and gaining deeper insights into the mechanisms of action of herbal medicines.

## 4. Discussion

Uncovering the synergistic mechanisms among herbal medicines is a significant challenge due to their complex components and interactions. Currently, most TCM databases provide only target information of ingredients linked to diseases through target-disease associations or used for pathway enrichment analysis. However, herb-disease associations arise from more intricate interactions than the additive effects of individual ingredients. To address these limitations, we developed HerbComb, a data portal that offers diverse herbal pharmacological data and supports combinatorial analysis and drug repurposing. HerbComb employs a network proximity-inference model to quantitatively characterize the interactions between herbs, ingredients, and diseases, thereby identifying their synergistic mechanisms with these key functions: 1) **Drug repurposing:** Systematic modeling to identify potential therapeutic and synergistic ingredients/herbs for specific diseases on a large scale. Notably, a combinatorial interaction network is provided to help users visually understand the synergistic mechanisms. 2) **Customized combinatorial analysis:** HerbComb enables users to perform tailored analyses of specific herb/ingredient combinations for diseases, simulating the decision-making process in herbal medicine practic. 3) **Biological and pharmacological features:** To further facilitate the combinational analysis, multiple biological and pharmacological features of herbal ingredients, including their gene signatures and ADMET properties, were also interrogated as interactive functionalities.

Our null distribution (comprising 304,992 randomly sampled ingredient pairs) approximates a normal distribution (**Figure 3B**). The 5% significance threshold was determined empirically from this distribution. Here, we intentionally included similar ingredient pairs in our random sampling because they reflect real-world TCM practice according to the “JUN-Chen-Zuo-Shi” theory, where similar herbs are combined to enhance therapeutic effects. Excluding them would artificially bias the null distribution against clinically relevant combinations. However, with over 50,000 ingredients, exhaustive pairwise analysis (approximately 1.25 × 10^9 unique pairs) is computationally prohibitive. Instead, we constructed our null distribution here using all ingredient pairs that co-occur in existing formulae (n = 304,992). This strategy enables us to capture the proper distribution of target set proximities in our database, which reflects real TCM practice while maintaining statistical validity against an appropriate null model and reducing computational requirements from 2.5 × 10^12^ to 10^5^ distance calculations. The extreme 5% tail (lowest network distances) identifies pairs that are significantly closer than 95% of random expectations. However, the network proximity approach identifies pairs with exceptionally close target interactions, a strong indicator of potential functional interplay (either synergistic or additive). However, it does not pharmacologically quantify the combined effect relative to individual effects, thereby not definitively distinguishing synergy from additivity. Furthermore, this method primarily focuses on identifying potential positive interactions (synergy/additivity) and does not specifically predict antagonistic interactions, which might involve opposing pathways not captured by simple distance minimization. Future work will incorporate antagonism detection through pathway opposition metrics. ?

While we have established a comprehensive platform for the combinatorial analysis of herbal medicines, several methodological constraints arising from data limitations warrant consideration. First, our approach inherently prioritizes well-studied herbs (e.g., *Glycyrrhiza uralensis, Panax ginseng*) with abundant target data, potentially overlooking under-investigated botanicals. This reflects systemic biases in existing literature and databases used for compilation. Consequently, network predictions may favor frequently studied herbs, potentially overlooking synergistic potential in understudied botanical resources. Similarly, our manually curated interaction dataset may overrepresent positive findings due to the preferential publication of synergistic results versus neutral/antagonistic outcomes. Third, high-confidence synergistic pairs often correspond to classical formulations (e.g., *Coptis chinensis* and *Scutellaria baicalensis* in Huang-Lian-Jie-Du-Tang). While validating traditional knowledge, this may limit the discovery of novel combinations that extend beyond established paradigms. Our network proximity approach partially mitigates this by evaluating biological relationships rather than co-prescription frequency.

HerbComb provides ADMET-aware screening (identifying 9 of 32 ischemic stroke-targeted compounds with favorable bioavailability and blood-brain barrier permeability in **Figure 3C**). However, these predictions serve only as initial pharmacokinetic filters. Although ADMET profiling offers valuable insights, true organism-level effects require consideration of additional biological complexities, including gut microbiome interactions (biotransformation, metabolic activation), tissue-specific distribution, and off-target signaling effects. More importantly, HerbComb is a hypothesis-generating tool. Thus, all predictions require experimental confirmation via in vitro, organoid, or in vivo studies. Additionally, traditional knowledge remains essential for contextualizing results. Future improvements will incorporate microbiome interaction predictions, dynamic PK/PD modeling, and clinical correlation data to enhance combination drug discovery.

In summary, we developed HerbComb, a novel web-based platform for characterizing synergistic interactions among herbal medicines. This platform provides new insights into synergistic mechanisms and facilitates the discovery of effective drug combinations for disease treatment.

## Acknowledgments

This work was supported by the Jiangsu Province Science Foundation for Youths (No. BK20231024), the Young Scientists Fund of the National Natural Science Foundation of China Grants (No. 82405199), the National Key Research and Development Program of China (No. 2022YFC3500201), and the Jiangsu Qinglan project (2024). It was also supported by funding from the Academy of Finland projects 317680 (J.T.) and 320131 (J.T.).

## Data and Software Availability

The datasets supporting this study are available upon request. For the review process, the data can be accessed via a restricted Zenodo private link: https://tinyurl.com/mwya586m. The data includes the ADMET properties of ingredients, as well as information on formulas, traditional Chinese medicines, ingredients, and related diseases.

## Notes

### Competing Interest Statement

The authors have declared no competing interest.

